# Whole-Genome Sequencing of the Wild Barley Diversity Collection: A Resource for Identifying and Exploiting Genetic Variation for Cultivated Barley Improvement

**DOI:** 10.1101/2024.11.18.624148

**Authors:** Ahmad H. Sallam, Yu Guo, Murukarthick Jayakodi, Axel Himmelbach, Anne Fiebig, Jamie Simmons, Gerit Bethke, Yoonjung Lee, Rebecca Spanner, Ana Badea, Michael Baum, François Belzile, Roi Ben-David, Robert Brueggeman, Austin Case, Luigi Cattivelli, Michael Davis, Christoph Dockter, Jaroslav Doležel, Antonin Dreiseitl, Ryan Gavin, Lior Glick, Stephan Greiner, Ruth Hamilton, Patrick M. Hayes, Scott Heisel, Cynthia Henson, Benjamin Kilian, Takao Komatsuda, Chengdao Li, Cheng Liu, Ramamurthy Mahalingam, Maren Maruschewski, Oadi Matny, Andreas Maurer, Klaus F. X. Mayer, Itay Mayrose, Peter Morrell, Matthew Moscou, Gary J. Muehlbauer, Youko Oono, Frank Ordon, Hakan Özkan, Ales Pecinka, Dragan Perovic, Klaus Pillen, Mohammad Pourkheirandish, Joanne Russell, Jan Šafář, Silvio Salvi, Miguel Sanchez-Garcia, Kazuhiro Sato, Thomas Schmutzer, Uwe Scholz, Jeness Scott, Gurcharn Singh Brar, Kevin P. Smith, Mark E. Sorrells, Manuel Spannagl, Nils Stein, Alessandro Tondelli, Roberto Tuberosa, James Tucker, Thomas Turkington, Jan Valkoun, Ramesh Pal Singh Verma, Marcus A. Vinje, Maria von Korff Schmising, Jason G. Walling, Robbie Waugh, Roger P. Wise, Brande B. H. Wulff, Shengming Yang, Guoping Zhang, Martin Mascher, Brian J. Steffenson

**Author notes:** **Corresponding author** Correspondence to Brian Steffenson and Martin Mascher.

## Abstract

To exploit allelic variation in *Hordeum vulgare* subsp. *spontaneum*, the Wild Barley Diversity Collection was evaluated for several agronomic traits and subjected to paired-end Illumina sequencing at ∼9X depth, generating 109.5 million single nucleotide polymorphisms after alignment to the Morex V3 assembly. A genome-wide association study of lemma color identified one marker-trait association (MTA) on chromosome 1HL close to *HvBlp*, the cloned gene controlling black lemma. Four MTAs were identified for stem rust resistance: one co-locating to the complex RMRL1-RMRL2 locus on 5HL, and three novel loci on 1HS, 1HL, and 5HL. Six MTAs for days to heading (DTH) on vernalized plants were identified on all chromosomes except 1H and 6H. Two MTAs for DTH on non-vernalized plants were identified on chromosomes 1HL and 2HS. All MTAs for DTH were novel. The whole genome sequence data described herein will facilitate the identification and utilization of new alleles for barley improvement.

## Background and Summary

Barley (*Hordeum vulgare* L. subsp. *vulgare*) was one of the first crops domesticated in the Near East about 10,000 years ago (Zohary et al. 2012) and today is cultivated over 47 million hectares worldwide (FAO 2017). Its main uses include malt for various alcoholic beverages, feed for animals, and food for humans. Through the domestication process and modern plant breeding, the genetic diversity of barley has been dramatically eroded, leaving the crop vulnerable to various biotic and abiotic threats and limiting further improvements for key traits. The primary gene pool of barley includes varieties, breeding lines, landraces, and wild barley (*H. vulgare* L. subsp. *spontaneum* C. Koch. Thell.), the latter of which can readily hybridize with the cultivated forms (Harlan and Zohary 1966; Liu et al. 2024). Studies aimed at identifying unexploited genes for use in barley breeding programs generally include panels that are more closely related to elite germplasm so as to not disrupt the genetic linkages of favorable alleles for yield, quality and agronomic traits. When a particular trait cannot be found in the cultivated forms of the primary gene pool, researchers often seek the desired alleles in the wild progenitor. To capture the allelic variation that exists in wild barley, an ecogeographically diverse collection (Wild Barley Diversity Collection; WBDC) was assembled (Steffenson et al. 2007). This collection is composed of 318 accessions from across the range of *H. v*. subsp. *spontaneum* and has been evaluated for various agronomic, morphological, nutritional, and disease/pest resistance traits. These evaluations revealed a high level of variation for nearly all of the characterized traits, leading to subsequent genetic and genome-wide association studies (GWAS) based on various molecular marker technologies (Roy et al. 2010; Sallam et al. 2017; Mahalingam et al. 2020; Xu et al. 2021; Walling et al. 2022). Here we describe the whole genome sequencing (WGS) of 281 WBDC accessions with ∼9X coverage and demonstrate its utility for identifying novel and previously described genes in *Hordeum vulgare* using an association genetic approach.

## Methods

### Wild barley germplasm

Collection site data for longitude and latitude, elevation, high and low temperature, rainfall, and soil type (**Table S1; Figure 1a**) were used to assemble the WBDC at the International Center for Agricultural Research in the Dry Areas (ICARDA) (Steffenson et al. 2007). The proportion of samples included were generally reflective of the density of populations in the Fertile Crescent, Central Asia, North Africa and Caucasus regions. Of the 318 originally selected WBDC accessions, 37 were dropped due to failed genotyping/sequencing, duplication or seed admixtures. The final sequenced panel is comprised of 281 accessions from 19 countries. Single plant selections were initially made from each accession and then selfed for five successive generations in the greenhouse before being used for DNA extraction and sequencing.

**Figure 1.**
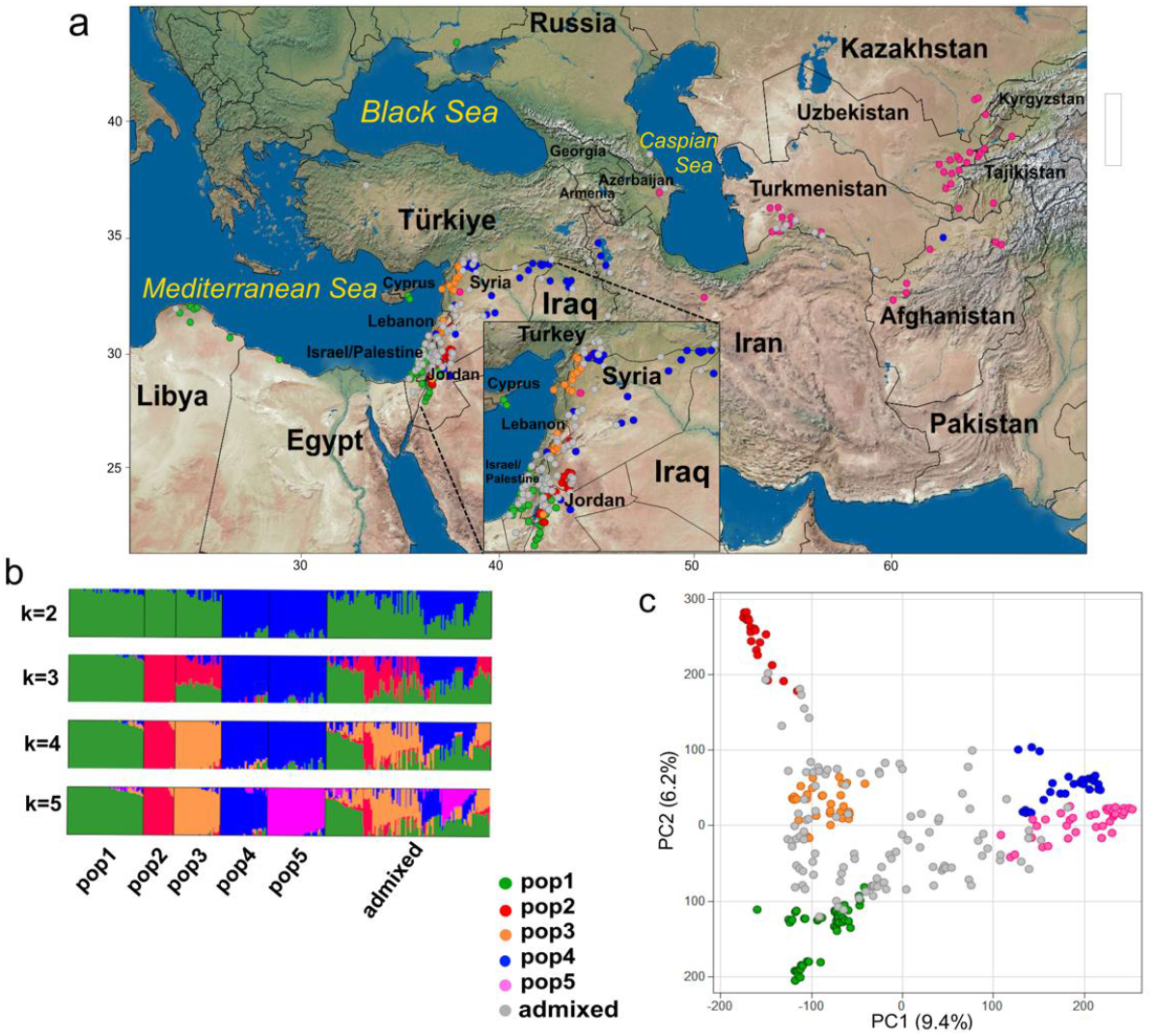
a) Geographic distribution of 281 *Hordeum vulgare* subsp. *spontaneum* accessions of the Wild Barley Diversity Collection (WBDC) and population designations based on whole genome sequencing; b) Population structure of the WBDC inferred from ADMIXTURE where the number of populations identified was five; and c) Principal component analysis determined from 571,285 SNPs where five major groups that corresponded closely with the population assignment results found with ADMIXTURE. Admixed accessions are shown in gray.

### DNA extractions

The first and second leaves of each accession were harvested, flash-frozen in liquid nitrogen, and stored at -80°C until the DNA extractions were performed. For the extractions, tissue was first ground to a fine powder in liquid nitrogen using a mortar and pestle. Then genomic DNA was extracted using a modified CTAB protocol (Yu et al. 2017). Agarose gel electrophoresis was used to confirm that the genomic DNA was of high molecular weight (>10 kb). DNA quality was assessed using a NanoDrop spectrophotometer.

### WGS library preparation and sequencing

WGS libraries were prepared using the ‘Illumina Nextera DNA Flex Library Preparation Kit’ (workflow for 100-500 ng DNA input, 5 PCR-cycles for the addition of indexes) according to manufacturer’s instructions (Illumina, Inc., San Diego, CA, USA). The final library pool was quantified by qPCR (Mascher et al. 2013). The pool was sequenced (XP workflow, paired-end, 2 × 151 cycles) using the Illumina NovaSeq6000 device and standard protocols from the manufacturer.

### Variant calling

Raw reads were trimmed with cutadapt (Martin 2011) and aligned to the Morex (V3) reference genome assembly (Mascher et al. 2021) using Minimap2 (Li 2018). Alignments were sorted using Novosort (http://www.novocraft.com). BCFtools (Danecek et al. 2021) was used to call single nucleotide polymorphisms (SNPs) and short indels.

### ADMIXTURE analysis

To analyze individual ancestry coefficients with ADMIXTURE (version 1.23) (Alexander et al. 2009) raw SNPs (marker set 1) were filtered by retaining bi-allelic sites with read depth ≥2 and ≤50 (otherwise set to missing), <20% missing calls, <20% heterozygous calls and minor allele frequency (MAF) ≥5%. Prior to ADMIXTURE runs, linkage disequilibrium pruning was done with PLINK (version 1.9) (Purcell et al. 2007) using the parameters “--indep-pairwise 50 10 0.1”. For each value of K (range of 2-5), 20 replicate runs of ADMIXTURE were combined with CLUMPP (version 1.1.2) (Jakobsson and Rosenberg 2007) and plotted with Distruct version 1.1 (Rosenberg 2004). The results of K are shown in **Figure 1b**. Individuals with ancestry of >80% from one inferred population are treated as non-admixed. The geographical distribution of the 281 WBDC accessions, together with their population structure (K=5) was plotted using JMP Pro software V17.0.0 (JMP software).

### Principal component analysis (PCA)

To select a robust and computationally manageable marker set for GWAS, several filtering steps were performed. Heterozygote calls were set to missing and bi-allelic sites with ≤10% missing data, ≥3% minor allele frequency and ≥ mean minimum read depth of 10 were retained (labeled marker set 2 with 1,414,415 SNPs) (**Table 1**). The SNP set was further pruned by discarding sites with *r*^2^ >0.2 in windows of 5 sites to generate the final marker set (marker set 3 with 571,285 SNPs) across the 281 WBDC accessions. PCA was performed on marker set 3 using the R-package GAPIT (Lipka et al. 2012).

**Table 1.** Number of single nucleotide polymorphism (SNP) markers identified from 281 *Hordeum vulgare* subsp. *spontaneum* accessions of the Wild Barley Diversity Collection after whole genome sequencing. Marker set 1 includes all identified SNPs after aligning with the Morex V3 assembly. Marker set 2 was derived from Marker set 1 after converting heterozygotes to missing data and retaining bi-allelic sites with ≤10% missing data, ≥3% minor allele frequency and ≥ mean minimum read depth of 10. Marker set 3 was generated by discarding sites with *r*^*2*^ > 0.2 in windows of 5 sites.

|  | Marker Set 1 | Marker Set 2 | Marker Set 3 |
| --- | --- | --- | --- |
| 1H | 13,593,327 | 179,422 | 75,920 |
| 2H | 16,919,413 | 208,999 | 79,219 |
| 3H | 17,346,279 | 219,784 | 88,245 |
| 4H | 16,291,194 | 234,082 | 101,368 |
| 5H | 14,546,251 | 175,214 | 69,269 |
| 6H | 13,588,000 | 174,716 | 67,441 |
| 7H | 17,214,958 | 222,198 | 89,823 |
| **Total** | 109,499,422 | 1,414,415 | 571,285 |

### Phenotyping of traits

To demonstrate the utility of the WGS dataset for positioning loci in wild barley, we selected three important traits for study: 1) lemma color, 2) stem rust resistance and 3) heading date under both vernalized and non-vernalized treatments. Lemma color was assessed by taking digital images of the seeds and then analyzing each color channel using the Fiji package (Schindelin et al. 2012) (**Supplemental Information**). Stem rust assays on seedling plants were performed with two races (MCCFC and QCCJB) of the wheat stem rust pathogen (*Puccinia graminis* f. sp. *tritici* (*Pgt*) and one isolate (92-MN-90) of the rye stem rust pathogen (*P. graminis* f. sp. *secalis*) (*Pgs*) as described in Sallam et al. (2017). Days to heading (DTH) was assessed on vernalized and non-vernalized plants in the greenhouse (**Supplemental Information)**.

### Genome-wide association mapping

To identify markers associated with the three traits, GWAS was conducted for 281 WBDC accessions using three different methods: 1) a Mixed Linear Model (MLM) that accounts for population structure (Q) + kinship (K) (Yu et al. 2006), 2) a Fixed and random model Circulating Probability Unification (FarmCPU) (Kusmec and Schnable 2018) that utilizes fixed and random effects iteratively to improve association power, and 3) a Bayesian-information and Linkage-disequilibrium Iteratively Nested Keyway (BLINK) (Huang et al. 2019) that utilizes Bayes and linkage disequilibrium to improve both association power and computation efficiency. All association mapping methods were executed in the R-package GAPIT. Marker trait associations (MTAs) identified with two or more methods or those detected with a single method but on two different datasets are presented. The Bonferroni test was performed to declare significant associations.

## Results and Discussion

The genomes of 281 WBDC accessions were sequenced with Illumina short reads, resulting in an average mapped read depth of ∼9X relative to the Morex V3 reference assembly (**Figure S1**). We discovered 109.5 million SNPs (**Table 1**) and 4.7 million short (1-50 bp) insertions and deletions (**Table S2, Figure S2**). Population structure was assessed by PCA and model-based ancestry estimation with ADMIXTURE using five ancestral populations (**Figures 1b,c**). Both yielded concordant results in that the populations defined by ADMIXTURE formed distinct clusters in PCA space. Consistent with prior results on the population structure of wild barley (Fang et al. 2014; Russell et al. 2016; Sallam et al. 2017), genetic relatedness mirrored geographic distance: the distribution of population centers roughly traced a path from the North African coast and the Southern Levant along the Fertile Crescent to Central Asia (**Figure 1a**). A detailed analysis of population structure in wild barley and its relationship to domesticated accessions using the present data set was undertaken by (Guo et al. 2025).

### Lemma color

Lemma color in the WBDC ranged from pale yellow (straw-colored) to brown and dark black based on visual inspection (**Figure 2**). Converted RGB values from digital images of pale yellow and dark black seed generally ranged from 10,750,000-13,550,000 and 4,350,000-6,570,000, respectively **(Table S1; Figure S3)**. Black lemma is a classic morphological trait in barley and is controlled by the *Blp* locus, which is composed of different alleles contributing to the intensity and distribution of color (Franckowiak and Lundqvist 1997). GWAS identified one association (WBDC_LC_1H_499.0) by a single SNP (S1H_499012052) on chromosome 1HL (**Table 2; Figure 3**). This SNP explained 17.3% of the phenotypic variation and lies in close proximity to *HvBlp*, the recently cloned gene controlling black lemma color positioned between 498.5 to 499.0 Mbp on 1H in the Morex (V3) assembly (Li et al. 2024). Due to the complexity of the locus and a duplicated fragment of *HvBlp*, it is difficult to state with certainty the physical relationship of the identified SNP marker and this gene.

**Table 2.**
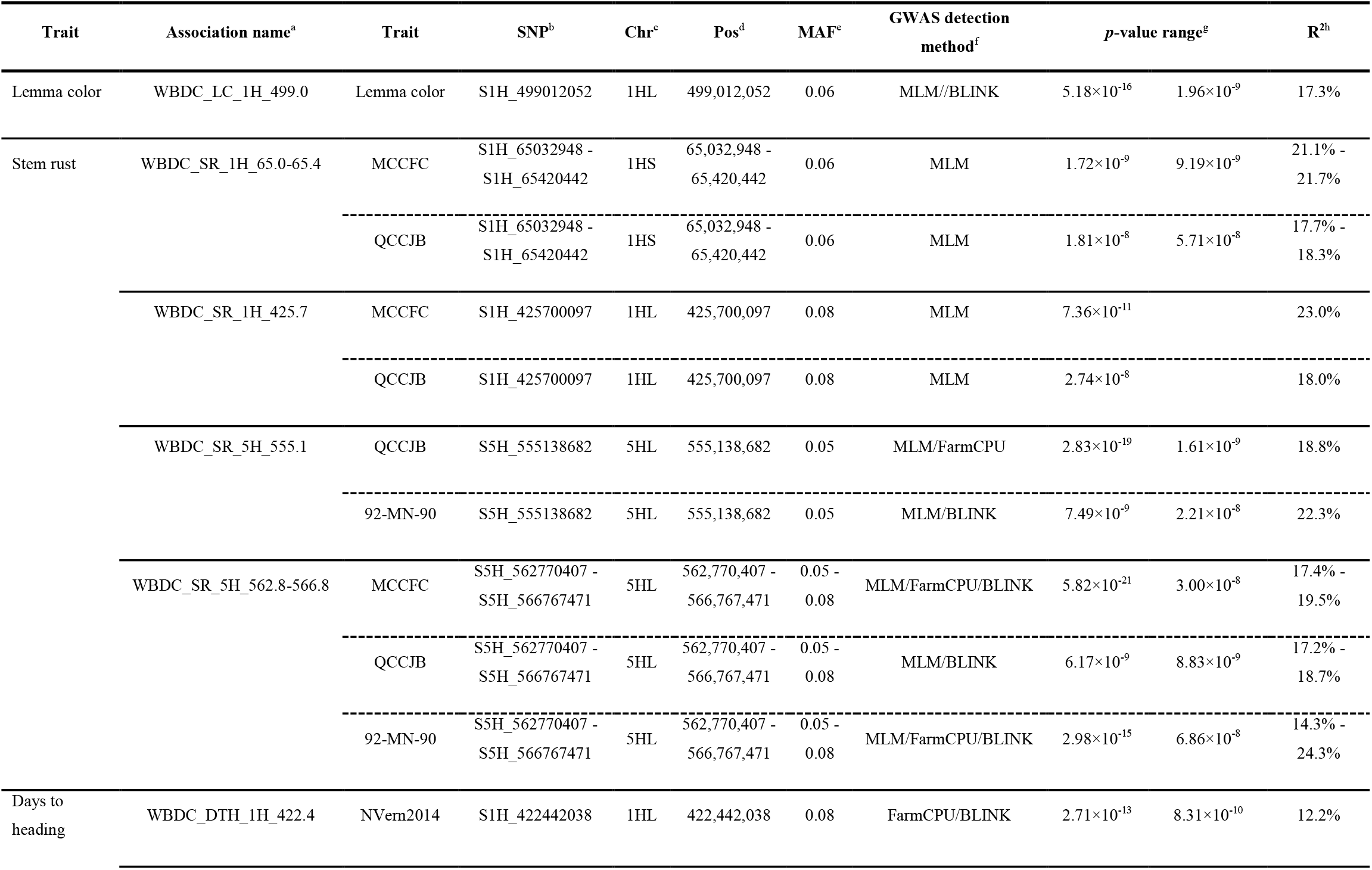
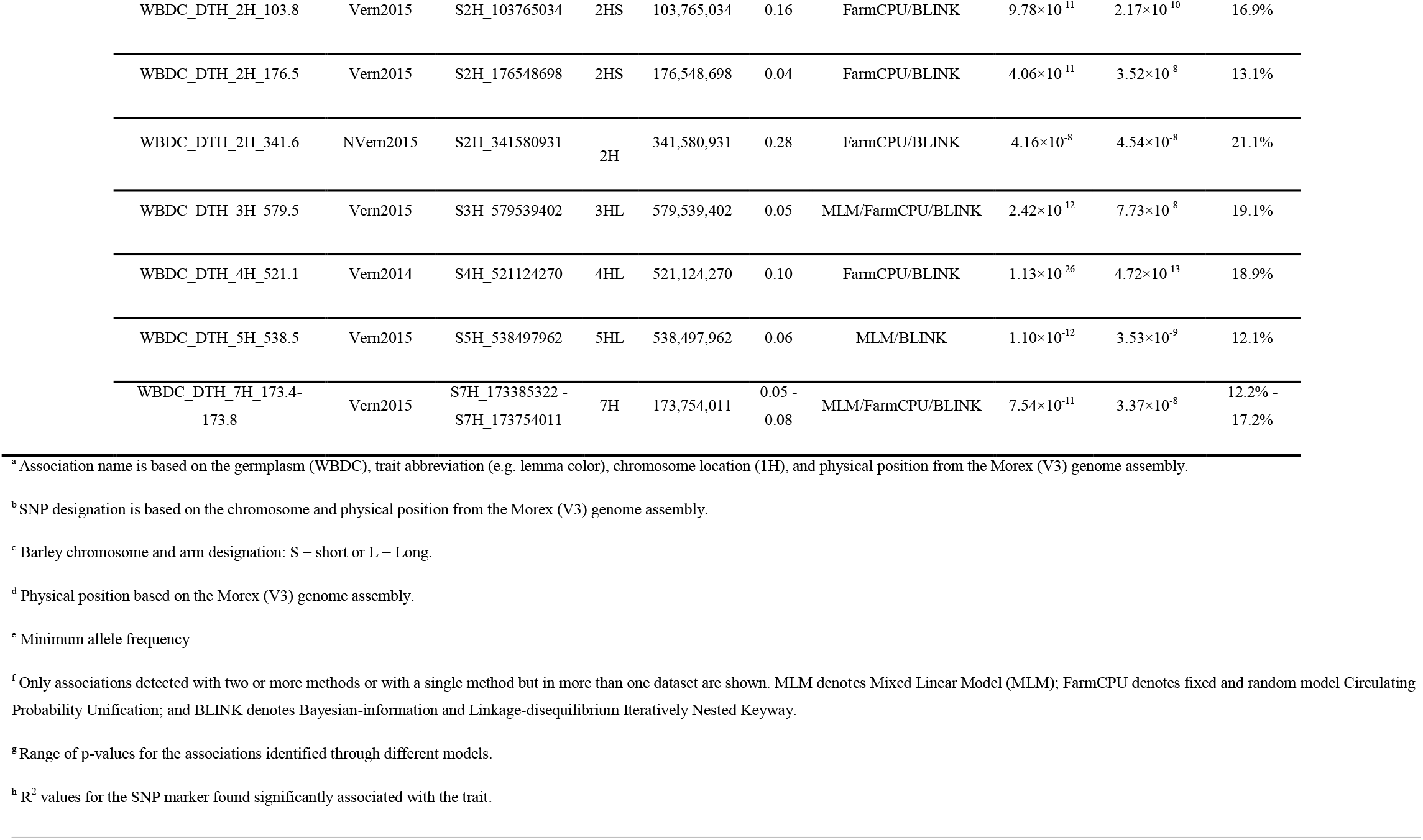
Single nucleotide polymorphism markers significantly associated with lemma color, stem rust resistance and days to heading in vernalized and non-vernalized plants in 281 *Hordeum vulgare* subsp. *spontaneum* accessions of the Wild Barley Diversity Collection.

**Figure 2.**
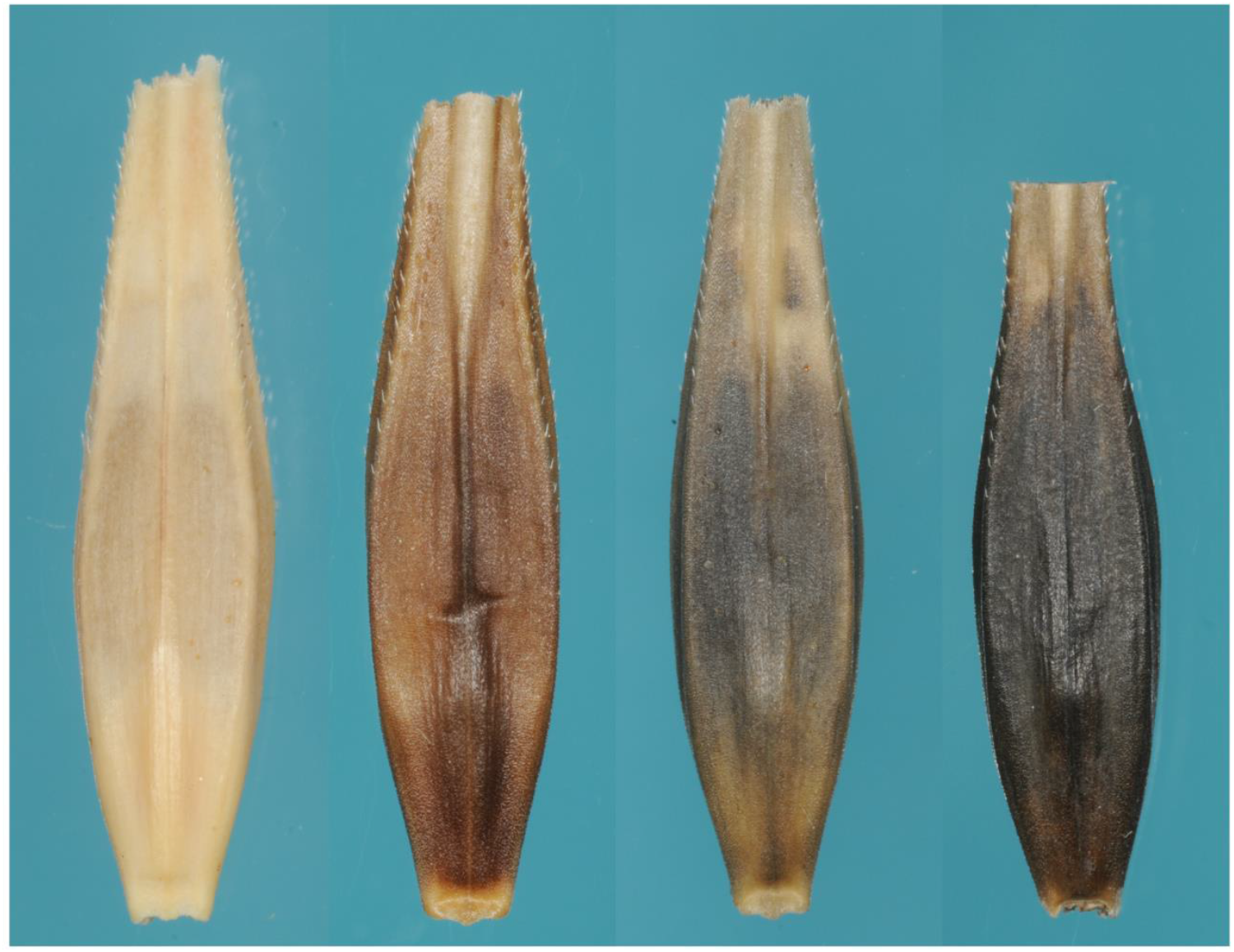
Examples of different lemma colors in the Wild Barley Diversity Collection; from left to right: yellow (straw) from WBDC045, brown from WBDC204, diffuse black from WBDC014, and dark black from WBDC355.

**Figure 3.**
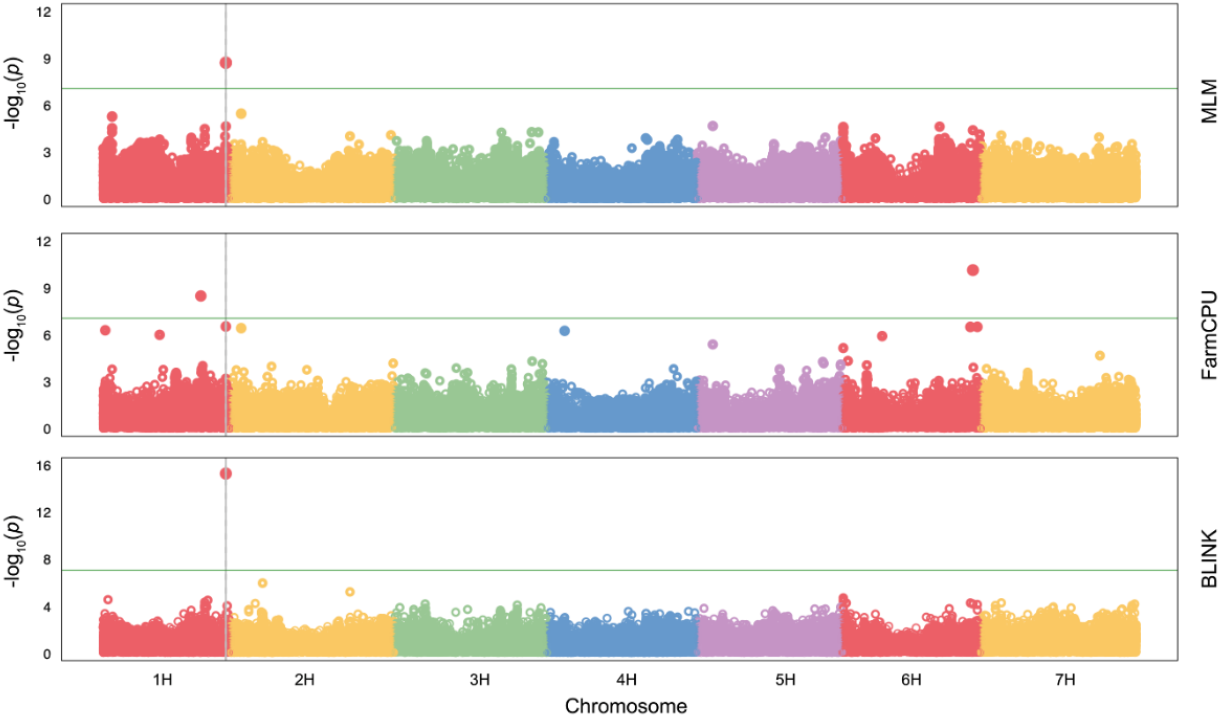
Manhattan plot displaying single nucleotide polymorphism (SNP) markers significantly associated with lemma color in the Wild Barley Diversity Collection. Three models were used in the analysis: 1) a Mixed Linear Model (MLM), 2) a Fixed and random model Circulating Probability Unification (FarmCPU), and 3) a Bayesian-information and Linkage-disequilibrium Iteratively Nested Keyway (BLINK). Bonferroni significance threshold is shown with a horizontal solid green line.

### Stem rust

Based on a coefficient of infection threshold of 2.7, only 15 (5.0%), 39 (14.0%), and 54 (19.0%) of the sequenced WBDC accessions were considered resistant to *Pgt*-MCCFC, *Pgt*-QCCJB and *Pgs*-92-MN-90, respectively (**Table S1; Figures S4a-c**). Four MTAs were identified for stem rust resistance to two or more cultures of *P. graminis* (**Table 2**; **Figures 4a-c**). Two novel MTAs were mapped on chromosome 1H in response to the *Pgt* races MCCFC and QCCJB. WBDC_SR_1H_65.0-65.4 and WBDC_SR_1H_425.7 were positioned on the short and long arms and explained 17.7-21.7% and 18.0-23.0% of the variation, respectively. WBDC_SR_5H_555.1 was mapped on chromosome 5HL in response to *Pgt*-QCCJB and *Pgs*-92-MN-90. This novel MTA explained 18.8-23.3% of the variation. WBDC_SR_5H_562.8-566.8 was identified on chromosome 5HL after challenge with all three *P. graminis* cultures and explained 14.3-24.3% of the variation. It co-located to the position of complex RMRL1-RMRL2 loci (Wang et al. 2013) from which several component resistance genes (e.g. *rpg4* and *Rpg5*) were cloned (Brueggeman et al. 2008; Arora et al. 2013).

**Figure 4.**
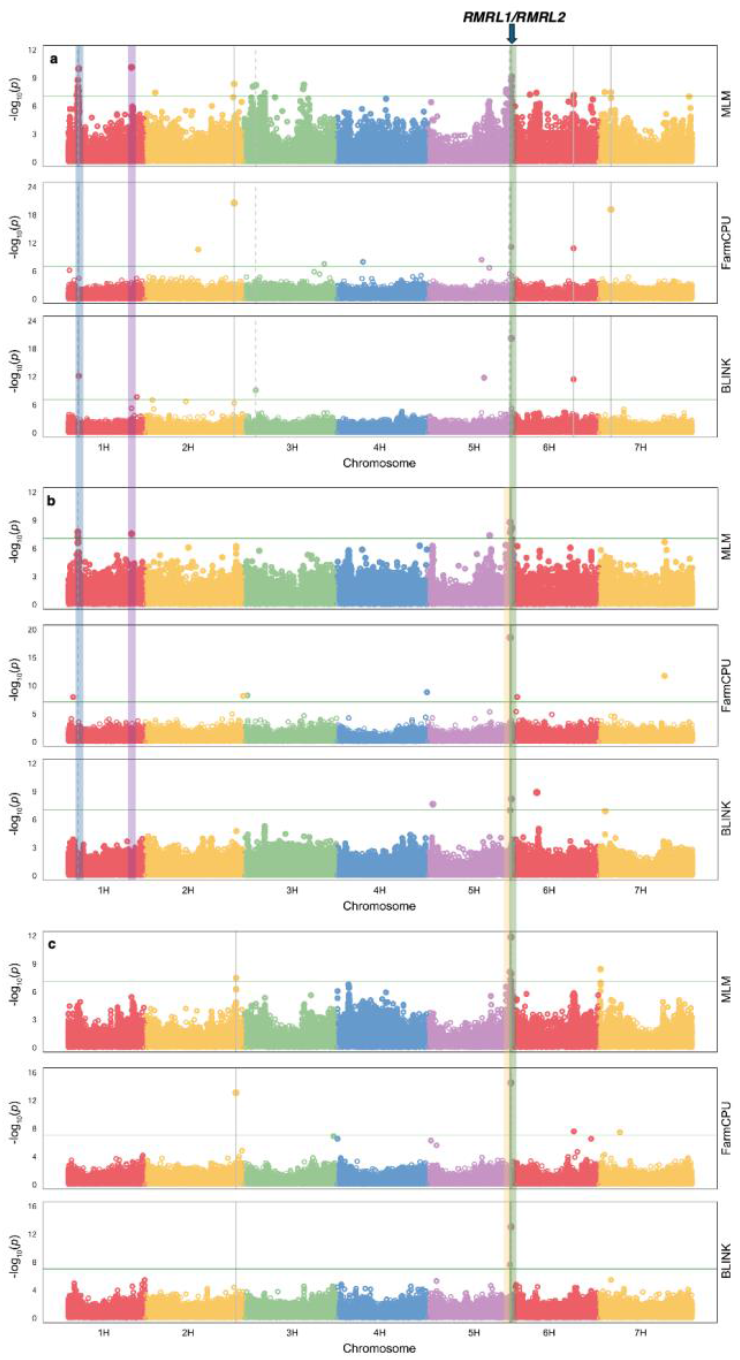
Manhattan plot displaying single nucleotide polymorphism (SNP) markers significantly associated with stem rust resistance to races (a) MCCFC and (b) QCCJB of the wheat stem rust pathogen (*Puccinia graminis* f. sp. *tritici*) and (c) isolate 92-MN-90 of the rye stem rust pathogen (*P. graminis* f. sp. *secalis*) in the Wild Barley Diversity Collection. Three models were used in the analysis: 1) a Mixed Linear Model (MLM), 2) a Fixed and random model Circulating Probability Unification (FarmCPU), and 3) a Bayesian-information and Linkage-disequilibrium Iteratively Nested Keyway (BLINK). Bonferroni significance threshold is shown with a horizontal solid green line. The vertical blue, purple, yellow and green lines show the significant associations consistently identified for a trait across two cultures of *P. graminis*. RMRL1/RMRL2 is a complex of several rust resistance genes.

### Days to heading under vernalized and non-vernalized treatments

A wide range of DTH was observed in the WBDC for both the vernalized and non-vernalized treatments. For the vernalized treatment, DTH ranged between 23-61 days and 34-89 days for the 2014 and 2015 experiments, respectively (**Table S1; Figure S5**). With respect to the non-vernalized treatment, DTH ranged between 20-145 and 42-147 days for the same respective years, with some accessions never heading after a growing period of over six months. Wild barley is regarded as a winter type cereal requiring a cold period to induce heading (Saisho et al. 2011). However, extended periods of growth can eventually induce heading as observed on accessions in the non-vernalized treatment (Muñoz-Amatriaín et al. 2020). High positive correlations were observed between the two datasets (2014 and 2015) for the vernalized (*r*^*2*^ = 0.69) treatment and for the non-vernalized (*r*^*2*^ = 0.71) treatment. In contrast, lower correlations (*r*^*2*^ = 0.19 to 0.30) were found between the datasets for the vernalized and non-vernalized plants, indicating the presence of different genes controlling heading date under these two treatments.

Six MTAs were identified for DTH in the vernalized treatment and two in the non-vernalized treatment (**Table 2; Figures 5a-d**). For the vernalized treatment, MTAs were identified on 2H (WBDC_DTH_2H_103.8 and WBDC_DTH_2H_176.5), 3H (WBDC_DTH_3H_579.5), 4H (WBDC_DTH_521.1), 5H (WBDC_DTH_5H_538.5) and 7H (WBDC_DTH_7H_173.4-173.8). The percentage of variation explained by these MTAs ranged from 12.1-19.1%. For the non-vernalized treatment, MTAs were identified on 1H (WBDC_DTH_1H_422.4) and 2H (WBDC_DTH_2H_341.6), explaining 12.2 and 21.1% of the variation, respectively. No common associations for DTH were identified between the different experiments. Additionally, all of the MTAs identified appear to be novel because they are not located near any of the previously described genes for heading date, including *Vrn-H1* and *Vrn-H2* on 5HL, *Vrn-H3* on 7HS, *Ppd-H1* on 2HS, or *Sdw1* on 3HL (Hu et al. 2019; Muñoz-Amatriaín et al. 2020; Sallam et al. 2024). The DTH genes identified in this study could be useful alternatives in breeding for heading date because of their large contribution to the variance and consistency of detection across GWAS methods. It is likely that these genes may have been lost during the domestication process and were therefore never incorporated into modern barley cultivars.

**Figure 5.**
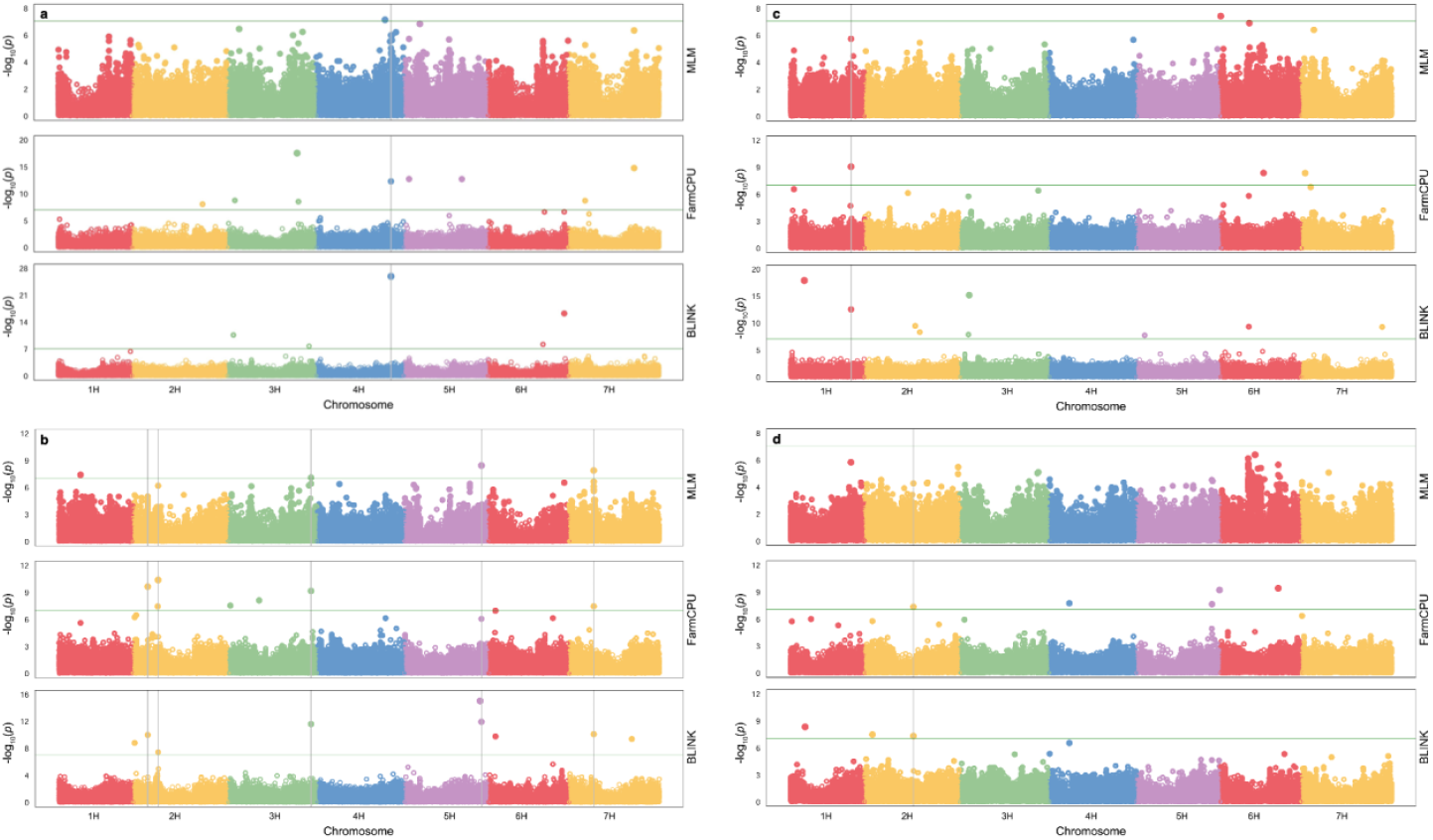
Manhattan plot displaying single nucleotide polymorphism (SNP) markers significantly associated with days to heading in vernalized accessions from the (a) 2014 experiment and (b) 2015 experiment and in non-vernalized accessions from the (c) 2014 experiment and (d) 2015 experiment of the Wild Barley Diversity Collection. Three models were used in the analysis: 1) a Mixed Linear Model (MLM), 2) a Fixed and random model Circulating Probability Unification (FarmCPU), and 3) a Bayesian-information and Linkage-disequilibrium Iteratively Nested Keyway (BLINK). Bonferroni significance threshold is shown with a horizontal solid green line.

WGS data for diverse accessions of a crop and its wild relatives are essential for population genomic studies, the informed selection of genotypes for full genome sequence assembly (pan-genomics) and the isolation of agronomically important genes. Our dataset complements similar short-read datasets for 1,315 domesticated barleys (Jayakodi et al. 2020; Jayakodi et al. 2024) and 100 wild barleys from another collection (Jayakodi et al. 2020). Chromosome-scale genome assemblies of nine WBDC accessions have been completed (Jayakodi et al. 2024), and we expect that more will follow in the future. Applying GWAS to the WBDC, we demonstrated the utility of high-coverage sequence data to identify novel genetic variation that may be useful in barley improvement. Additionally, we also validated major genes controlling key traits in barley such as *Blp* for black lemma color and RMRL1/RMRL2 for stem rust resistance. We anticipate that our dataset will also serve as a starting point for the identification of candidate genes underlying other important traits. In a companion paper, Guo et al. (2025) demonstrated the utility of WBDC sequence data in population genomic studies by analyzing them along with data for other diverse wild and domesticated barleys to reconstruct the evolutionary history of wild barley and elucidate the origin of haplotypes in cultivated barley. The sequenced genomes of the WBDC will help connect target phenotypic traits to chromosome positions. Reference genome positions, as identified by HORVU I.D.s in the Morex V3 assembly (Mascher et al. 2021), serve as anchors to protein-protein interactome hubs (Velásquez-Zapata et al. 2022) and the potential for engineering the molecular and cellular mechanisms by which key phenotypes are expressed.

### Web resources

Not applicable.

### Germplasm and Data availability

Seed of the complete WBDC (N=318) can be obtained from the USDA-ARS National Small Grains Collection as accessions PI 681726 to PI 682043. Raw sequence data are deposited in the European Nucleotide Archive (ENA) under project ID PRJEB56087. The variant data for marker sets 1, 2, and 3 (Table 1) have been deposited in the European Variation Archive (EVA) at EMBL-EBI under accession number PRJEB80165 (https://www.ebi.ac.uk/eva/?eva-study=PRJEB80165). Scripts used for variant calling, variant filtering and ADMIXTURE analyses are available in GitHub repositories at: https://github.com/SteffensonLab. We used stem rust reaction type data from a previously published G3 paper (Sallam et al. 2017): https://doi.org/10.1534/g3.117.300222. These data are also included in Table S1.

## Acknowledgments

A.H. acknowledges Ines Walde for expert technical assistance in the sequencing operation. B.J.S. thanks Harold Bockelman, Matthew Martin and Tamas Szinyei for technical assistance in the handling of the wild barley materials.

## Supplemental data

Supplementary Information, Figures S1-S5 and Tables S1-S2 are available to download as part of this submission.

## Usage notes

Not applicable

## Authors and Affiliations

As given above

## Contributions

B. J. Steffenson and M. Mascher designed and led the study. A. H. Sallam, G. Bethke, and J. Simmons grew and harvested the plant materials and extracted DNA. A. Himmelbach, and A. Fiebig performed the sequencing operations. A. H. Sallam, Y. Lee, M. Jayakodi, M. Mascher, R. Spanner, and Y. Guo analyzed the data. B. J. Steffenson, A. H. Sallam, Y. Lee, R. Spanner, M. Mascher, and Y. Guo drafted manuscript and made the revisions. All authors reviewed and edited the manuscript.

## Corresponding author

Correspondence to Brian Steffenson and Martin Mascher

## Ethics declarations

Publisher’s note Springer Nature remains neutral with regard to jurisdictional claims in published maps and institutional affiliations.

